# Encoding of movement primitives and body posture through distributed proprioception in walking and climbing insects

**DOI:** 10.1101/2024.09.27.615364

**Authors:** Thomas van der Veen, Volker Dürr, Elisabetta Chicca

## Abstract

Targeted reaching movements and spatial coordination of footfall patterns are prime examples of spatial coordination of limbs in insects. To explain this, both physiological and computational studies have suggested the use of movement primitives or the existence of an internal body representation, much like they are assumed to occur in vertebrates. Since insects lack a dedicated posture-sensing organ or vestibular system, it is hypothesized that they derive high-level postural information from low-level proprioceptive cues, integrated across their limbs. The present study tests the extent to which a multi-layer spiking neural network can extract high-level information about limb movement and whole-body posture from information provided by distributed local proprioceptors. In a preceding part of the study, we introduced the phasic-tonic encoding of joint angles by strictly local proprioceptive hair field afferents, as well as high-accuracy encoding of joint angles and angular velocities in first-order interneurons. Here, we extend this model by second-order interneurons that use coincidence detection from two or three leg-local inputs to encode movement primitives of a single leg. Using experimental data on whole-body kinematics of unrestrained walking and climbing stick insects, we show that these movement primitives can be used to signal particular step cycle phases, but also step cycle transitions such as leg lift-off. Additionally, third-order interneurons are introduced to indicate climbing behaviour, for example by encoding the body pitch angle from 6 *×* 3 local leg joints. All encoding properties are validated against annotated experimental data, allowing for relevance rating of particular leg types and/or leg joint actions for all measures encoded. Our results demonstrate that simple combinations of two or three position/velocity inputs from disjunct hair field arrays are sufficient to encode high-order movement information about step cycle phases. The resulting movement primitive encoding may converge to represent particular locomotor states and whole-body posture.

**Author summary:** Insect behaviours such as navigation or climbing involve complex movement sequences that have led scientists to postulate the existence of an internal body representation. As insects lack a dedicated organ for monitoring body posture, a major problem in computational neuroscience and biomimetic robotics is how high-level information about body posture and coordinated movement may be extracted from distributed, local, low-level sensory measures, such as joint angles or angular velocities. To solve this problem, we developed a spiking neural network model. The model was tuned and evaluated with experimental data on complex climbing sequences of stick insects, with detailed information about 6 *×* 3 joint angle time courses. In a preceding study, we focused on how joint angle sensors encode this information at various body parts and how it is processed to represent local joint position and movement. Here, we extend the model to include neurons that signal particular phases of a leg’s movement cycle. Other neurons encode whole-body movement, using the body pitch angle as an example parameter. We show that a straight-forward combination of movement signals from various body parts can indicate the timing of particular step cycle events, as well as provide an internal representation of the full body’s posture.

## Introduction

For locomotion in natural environments, animals require an elaborate sense of body size, shape, and posture, including reliable information on the spatial whereabouts of all limbs and appendages. In insects, experimental climbing paradigms have shown to involve distance estimates based on touch [1] or vision [2], with an essential component of the motor behaviour being aimed reaching for visual [3, 4] or tactile targets [5, 6]. Targeted limb movements also occur during grooming of the body surface [7–9] and in inter-leg coordination [10, 11]. Quite generally, spatial coordination problems solved in complex animal locomotion are believed to require short-term memory of spatial coordinates [12], mapping of limb postures [13, 14], and learning of the underlying sensorimotor transfer properties [15, 16] or internal representation of body size [17–19]. Our understanding of any of these physiological phenomena would greatly benefit from a computational model of distributed proprioception of limb posture and body posture [20, 21].

Given the striking absence of dedicated posture measurement organs such as statocysts, as found in some aquatic invertebrates like mollusks [22], or the vestibular system found in humans [23], insects need to estimate high-level posture information from various proprioceptors distributed across the animal’s body [24, 25]. Here, we propose a multi-layer spiking neural network (SNN) for encoding of coordinated movement of single legs and convergent information about whole-body posture. Building on a companion paper that modelled distributed afferent encoding of joint angles, and the extraction of high-accuracy position and velocity estimates in first-order interneurons (INs) (companion paper, van der Veen et al. [26]), the present paper expands the model by second-order INs encoding movement primitives of single limbs, and third-order INs encoding body pitch from 6 *×* 3 leg joints across the body. We use SNNs because they capture the temporal dynamics of biological neural systems. By encoding information in the precise timing of discrete spikes and incorporating neuron and synaptic states, SNNs are able to capture temporal patterns and exhibit great biological accuracy [27]. For a recent review of this topic, see Yamazaki et al. (2022) [28].

From a computational point of view, increasing levels of functional complexity in motor control differ in their numbers of Degrees of Freedom (DoF), and the associated redundancy [29, 30]. In the control of redundant motor systems, complexity is typically reduced by combining multiple DoF for synergistic action of several muscles, joint torques, or movement components [31, 32]. The first objective of the present paper is to apply the concept of movement primitives by establishing a pool of INs that signal posture-and-movement combinations of single legs, and by evaluating their specificity for encoding particular events throughout the step cycle. Locomotion is characterized by distinct states of the step cycle: the swing and stance phases. These states are mutually exclusive in that a leg cannot be in both phases simultaneously [33]. As these step cycle phases correspond to distinct states of mechanical coupling [34], several models of adaptive locomotion use distinct control modes for swing and stance [35, 36]. In cats [37] and locusts [38], INs have been identified that are activated during the swing or stance phases, or during the transition between them [38]. These INs could signal the control mode of the locomotor cycle that the leg is currently engaged in. Here we test whether an insect central nervous system (CNS) could use proprioceptive information about movement primitives to signal particular events or episodes of the step cycle.

The second objective of this paper is to test whether it is possible to extract high-level information about body posture from distributed, low-level sensory afferents. The question of how circuits at different levels of the CNS represent the body [21] remains an area that has been relatively under-researched. To date, computational models have enabled researchers to postulate hypotheses on the structure of neuronal pathways, with models addressing different levels of insect locomotion [39, 40], interlimb coordination [40, 41], descending interneurons [42], or insect proprioceptors [43, 44]. Here, we implement a layer of third-order INs to test whether distributed, concurrent encoding of movement primitives from six legs is sufficient to tell the behavioural states ‘climbing’ from ‘level walking’. In fact, proprioceptive afferent activity from hair plates has been shown to be sufficient to estimate the body pitch of a climbing stick insect using a multi-layered artificial neural network [45]. Expanding on this knowledge, our present model uses incremental encoding of increasingly complex information about whole-body kinematics to encode body pitch in the spike rate of third-order INs. All SNN layers of our model are optimized or evaluated with experimental data on whole-body kinematics of unrestrained walking and climbing in the stick insect *Carausius morosus*.

The manuscript is structured as follows: The Methods section introduces the dataset and provides a detailed explanation of the extended SNN architecture, describing the methodology layer by layer. The results section then presents our findings for each network layer. Finally, the discussion concludes with a comparison of modeling results and physiological evidence and a general outlook on future research on distributed proprioception.

## Methods

### Dataset

The experimental data used in this study comprises complete body kinematics from unrestrained walking and climbing stick insects. It was initially collected to compare the whole-body kinematics of walking and climbing among closely related stick insect species with different body morphologies [46] and for characterization of distinct step classes [47]. The data set is the same as that used by [45] and the companion paper (van der Veen et al., companion paper [26]). Nine specimens of the species *Carausius morosus* (de Sinéty, 1901) walked freely on a horizontal walkway that had a width of 40 mm and a length of 490 mm. The animals encountered either a flat surface (not shown, 78 trials), or two steps of 48 mm in height (Fig 1ai, 15 trials). In the trials used here, stick insects climbed two stairs of 48 mm height. This height exceeded the leg length and was equivalent to three times the body clearance.

**Fig 1.**
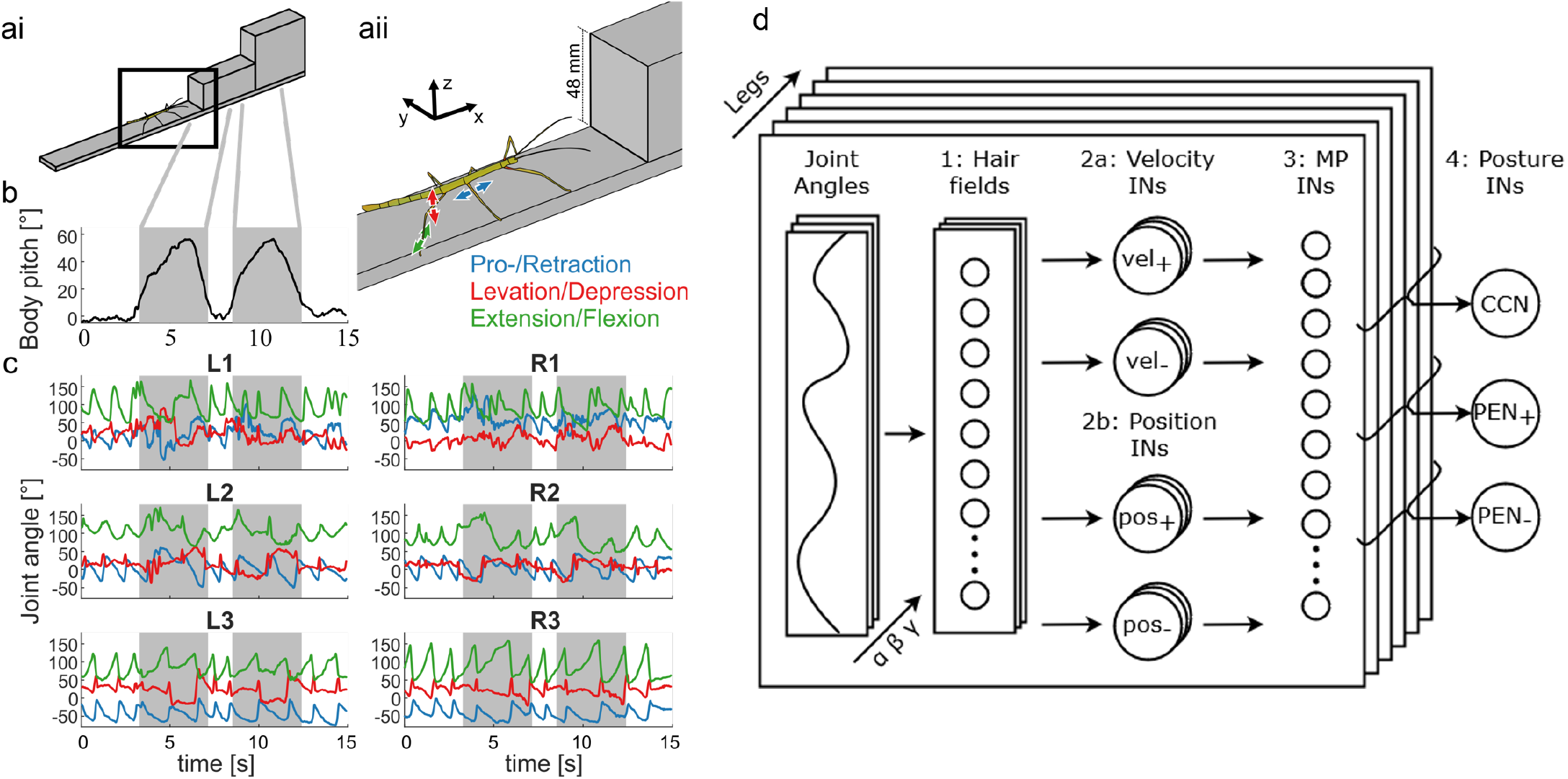
Experimental setup and SNN architecture. Setup (a), body pitch (b) and joint angle time courses (c) of a single trial. ai: Whole setup with a stick insect in front of the first stair. The black box indicates the enlarged area in aii. aii: Blue, red and green arrows indicate movement directions for pro-/retraction, levation/depression and extension/flexion, respectively. b: Pitch angle of the body axis over time of a representative trial. Grey shading in b and c indicates a body pitch above 10° in which the animal can be considered as climbing. c: joint angles of the same trial as in b. The six panels correspond to the six legs of the animal, with front legs, middle legs and hind legs in the top, middle and bottom panels, respectively. Left legs on the left and right legs on the right three panels. The colour code of the three joint angles per leg matches the movement directions explained in aii. d: SNN architecture. The architecture from the companion paper (van der Veen et al., companion paper [26]) is extended by 112 movement primitive (MP) INs per leg. These leg-specific neurons fire whenever combinations of joint angles or movements occur, making them indicators of particular phases of the step cycle or during phase transitions. In the fourth layer, three types of posture INs collect converging information from MP INs of all six legs. During climbing (shaded region in b), the climbing classifier neuron (CCN) spikes to indicate a ‘state of climbing’, and two pitch estimator neurons (PENs) provide a time-resolved estimate of the body pitch angle.

*C. morosus* has six legs with similar morphological structure, i.e. without obvious functional differences among them. Each leg comprises a short basal coxa, a fused trochantero-femur, a long and thin tibia, and a distal tarsus with five tarsomeres. The dataset includes motion capture data on three joint angles per leg. Arrows in Fig 1aii show the movement directions of these pivotal joints: the thorax-coxa joint facilitates protraction-retraction movements (blue); the coxa-trochanter joint allows for levation-depression movements (red); and the femur-tibia joint governs flexion-extension movements (green). Throughout this work, these three joints will be referred to as the *α, β*, and *γ* joints, respectively. The front, middle, and hind legs are denoted as 1, 2, and 3, respectively, for the right (R) and left (L) sides (Fig 1c).

The dataset was acquired with a marker-based motion capture system (Vicon MX10), comprising eight infrared cameras operating at 200 frames per second. Tracked markers were attached to the head, thorax, and all six legs of the insect. The captured marker trajectories were used to reconstruct the time courses of the three mentioned leg joint angles *α, β* and *γ*. A representation of the joint angles for each leg of a typical trial is shown in Fig 1c. For further details on the experimental procedure and data processing see [47].

### Spiking neural network (SNN) architecture

A schematic of the proposed SNN architecture is shown in Fig 1d. The two-layered architecture used in the companion paper (van der Veen et al., companion paper [26]), which included a sensory layer of proprioceptive hair field afferents and a layer of first-order INs, was extended by two further IN layers: First, a MP layer was added, consisting of 112 MP INs per leg. Each MP IN received spike trains from two or three position and velocity INs of the previous layer within the same leg, with each MP IN representing a unique combination of converging inputs, such that all possible configurations were covered. The MP INs functioned as coincidence detectors that triggered responses whenever their converging presynaptic inputs aligned temporally. The goal of the MP layer was to encode coordinated motion by two or three leg joints, thus capturing particular motion episodes of a single leg, such as step cycle phases (e.g., swing or stance).

Spike trains from all 6 *×* 112 MP INs, i.e., from all six legs, converged at the posture layer (i.e., layer four), which consisted of three distinct neurons. The purpose of the CCN was to signal the behavioural state of climbing, as opposed to regular walking. In contrast, the purpose of the PENs was to test whether a reliable estimate of body pitch angle could be obtained from distributed proprioception of limb kinematics. To do so, *PEN*_+_ and *PEN*_−_ were tuned to fire during climbing and regular walking, respectively, with their difference providing a time course for body pitch.

Both the MP INs and posture INs were modeled by a simplified leaky integrate-and-fire (LIF) model, as it provided a linear relation between input spike rate and output spike rate with a leakage term. The proposed architecture was implemented directly in Python version 3.9. The simulations were conducted on a system equipped with 16 GB of RAM and an AMD Ryzen 5600x processor. The differential equations that describe the behaviour of spiking neurons and synapses were solved over time using the backward difference method, using a time step of *dt* = 0.25 ms.

### Leaky integrate-and-fire (LIF) model

Due to its integration capabilities, simplicity, and small parameter set, the LIF model was selected for all neurons modeled in this work [48]. In this model, a pre-synaptic spike that occurs at time *t*^pre^ increases the membrane voltage *V* (*t*) by a synaptic weight *ω* [49]:

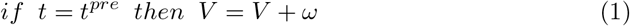

The dynamics of the LIF are as follows:

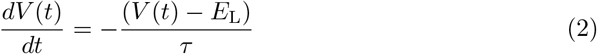

where *τ* is the decay time constant and *E*_L_ is the leak reversal potential. If the voltage *V* (*t*) exceeds the threshold voltage *V*_T_, a spike time *t*^*f*^ is recorded at that time step and the voltage is reset to *E*_L_:

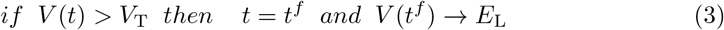

Given standard values for resting potential, *E*_L_, and spike threshold, *V*_T_, the only tunable parameters are *τ* and *ω*. For reference, the response of the LIF model to a constant frequency spike train stimulus is provided in S1 Fig.

### Layer three: Movement primitive layer

Layer two comprised two position INs and two velocity INs per joint, amounting to 12 INs per leg. The concept for layer three was to integrate converging input from two or three of these 12 position or velocity INs per leg (Fig 1d). Each one of the resulting 112 input combinations was to signal a particular type of motion and/or a particular joint angle domain. By doing so, the second-order INs in the MP layer acted as coincidence detectors, ensuring activation only when there was temporal alignment between presynaptic spikes in two or three first-order INs in the second layer. The MP layer was optimized using 78 trials of stick insects unrestrained walking without encountering steps.

#### Synapse connections

Each MP IN was modeled by the LIF model and was supplied with spike trains from two or three joints per leg. Specifically, each joint contributed information about either position or velocity, or none, with a synaptic connection from one of the joint’s four *pos*_+_, *pos*_−_, *vel*_+_, or *vel*_−_ INs, or no connection. Accordingly, each MP IN was described by a tuple of three entries - one entry per leg joint - allowing for any combination drawn from three sets of five. For example, the set for joint *α* was: 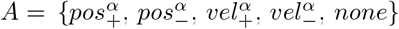, with analogous sets for joints *β* and *γ* as *B* and *C*, respectively. The possible combinations were obtained with a 3-fold Cartesian product (denoted by *×*) of the sets *A, B*, and *C*. This resulted in a set of ordered tuples (*a, b, c*), where *a, b*, and *c* are in *A, B*, and *C* respectively, expressed in set builder notation [50]:

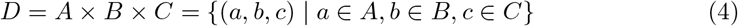

From the resulting set of 5^3^ = 125 ordered tuples, tuples with less than two active connections were invalid. Therefore, tuples with more than one *none* were excluded, resulting in a final count of 112 ordered tuples in set D. The set D naturally groups into seven subsets:

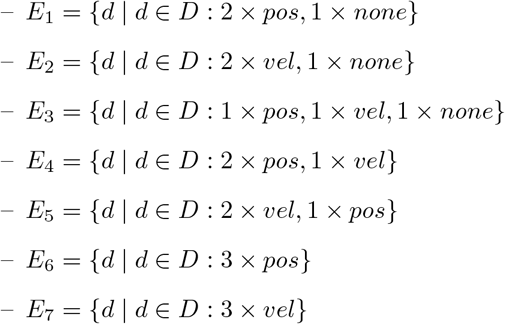

where *pos* represents either *pos*_−_ or *pos*_+_, and *vel* represents either *vel*_−_ or *vel*_+_. In each ordered tuple, the first element corresponded to the contribution of joint *α*, the second to joint *β*, and so forth. Therefore, the angle superscripts (e.g. ()^*α*^) were omitted. As an example, the subset *E*_3_ included the ordered tuple (*none, pos*_−_, *vel*_+_), where joint *α* had no contribution, joint *β* contributed spikes from the *pos*_−_ IN, and joint *γ* contributed spikes from the *pos*_+_ IN. In the remainder of the work, the subsets *E*_1_ through *E*_7_ are represented by: p-p, v-v, p-v, p-p-v, v-v-p, p-p-p and v-v-v, respectively. For each ordered tuple, exactly one corresponding MP IN had the specified presynaptic input. Therefore, there were 672 MP INs in total, with 112 second-order INs for each leg.

#### Evaluation of encoding performance

In order to evaluate the performance of MP INs, the ground truth reference was established based on the experimental data (actual condition), i.e., the real joint angles. To this end, reference spike trains of *pos*_+_ and *vel*_+_ INs were constructed by setting the element of the spike train to 1, denoting a spike, if the joint was in the positive domain or moving in the forward direction at time *t*, respectively. The element was set to 0, denoting no spike, if the real joint angle was in the negative domain or moving backward, respectively. Analogous spike trains were constructed for *pos*_−_ and *vel*_−_. Subsequently, 672 MP ground truth spike trains were constructed for the subsets *E*_1_ through *E*_7_, with the element set to 1 if spike trains coincided for all three ground truth inputs.

All of these 672 MP ground truth spike trains were then divided into 100 uniformly spaced bins. For each MP ground truth spike train, a bin was labeled as Positive (P) if the neuron spiked at least once during that binning period, and Negative (N) if no spike occurred. For comparison, all spike trains generated by MP INs (predicted condition) were binned in the same manner, labeled as predicted positive (PP) if a spike occurred at least once during that binned period, or predicted negative (PN) if zero spikes occurred.

The predicted and actual conditions were compared by constructing a confusion matrix, counting the number of true positives (TPs), true negatives (TNs), false positive (FPs) and false negatives (FNs). Due to the expected imbalance between P and N occurrences, simple accuracy calculations were deemed unreliable. Instead, the Matthew’s correlation coefficient (MCC) was chosen as a more suitable metric, a measure of the quality of binary (two-class) classifications [51]. Chicco and Jurman [52] demonstrated that the MCC was a more reliable statistical measure than the F1 score and accuracy, producing a high score only if the prediction achieved good results in all four categories of the confusion matrix (TP, FN, TN and FP). Its interpretation is similar to Pearson’s correlation coefficient: −1 indicates total negative correlation, 0 indicates no correlation, and 1 indicates total positive correlation [53]. The MCC is defined as [51]:

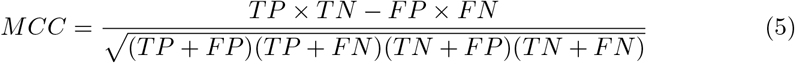

#### Evaluation of step cycle encoding

To evaluate how well an MP IN encoded a particular phase of the step cycle or, potentially, transitions between step cycle phases, the experimental data was first annotated according to whether a leg was in swing phase or rather in stance phase. A leg was annotated to be in swing phase whenever it was not in contact with the ground, whereas it was annotated to be in stance phase whenever it was in contact with the ground. Next, swing and stance phases were binned into eight equally sized bins each. Finally, for each MP IN, we determined the likelihood of a spike to occur in a particular bin, for all stance and swing phases of all 78 trials. Combining the results for the swing and stance phases yielded a phase vector 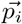 for MP IN *i*. Each one of these vectors consisted of 16 elements, where *p*_*i*,1:8_ represented the likelihood of spiking during the swing phase bins, and *p*_*i*,9:16_ represented the likelihood of spiking during the stance phase bins. If *p*_*i,j*_ was equal to 1, this indicated that the neuron consistently fired during that specific bin, whereas a value of 0 indicated no firing in that bin.

The phase vectors were then compared with target vectors corresponding to the swing phase 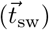 or the transition from swing to stance 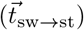 For instance, the target vector for the swing phase, 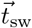of size 16, was defined such that the first 8 elements were set to 1 and the last 8 elements were set to 0. This target vector would perfectly align with the activity of a MP IN that fired at least once during each swing bin in the dataset, but never during any stance bin. The two target vectors are shown below, arranged visually in two rows. The top row corresponds to the swing bins, and the bottom row corresponds to the stance bins:

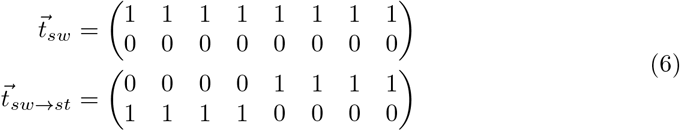

Encoding of particular step cycle phases by the *i*-th MP IN was then evaluated by means of score *S*_phase,*i*_ for either one of the two target vectors. For this we used the difference between the target and phase vectors as follows:

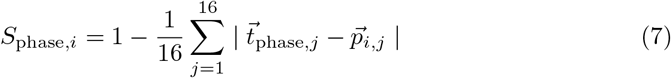

Here,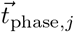represents the *j*-th element of the target vector for the given phase, and 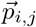 represents the *j*-th element of the phase vector for MP IN *i*. This score quantified the deviation of the observed spiking activity from the idealized spiking pattern for the specified phase, where a score of 1 indicated perfect alignment and 0 anti-alignment. Note that target vector 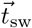 served to evaluate the encoding of both swing and stance because swing and stance were mutually exclusive states and, therefore, being in swing phase equaled NOT being in stance phase, and vice versa. The same held for target vector 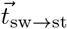 which served to evaluate the encoding of both the swing-to-stance transition (i.e., touch-down) and the stance-to-swing condition (i.e., lift-off).

### Layer four: Posture layer

With 672 unique MP INs in layer three of the SNN, layer four was designed to read out this information in order to estimate a higher-order body parameter that required information from all six legs. Specifically our goal was to optimise encoding of body pitch. To achieve this, two PENs were connected to specific presynaptic neurons of layer three (Fig 1d), based on their spiking characteristics. The two PENs were to be tuned to spike during periods of high and low body pitch, respectively. The z-normalized difference in the spike rates of the PENs then served as an estimate for body pitch. As a complement, a single CCN was tuned to spike during climbing, i.e. when body pitch exceeded 10°.

#### Walking and climbing biases

For differential encoding of walking and climbing behaviour, the neurons in the MP layer (layer three) were tested for spiking biases for low or high body pitch angles, respectively. Specifically, all spikes of the primitive neuron *i* of leg *j* were counted as a ‘climbing spike’ 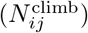if body pitch exceeded 10° at the time of this spike, or as a ‘walking spike’ 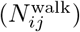 if body pitch remained below 10°. The ratio between ‘climbing spikes’ and ‘walking spikes’ was defined as follows:

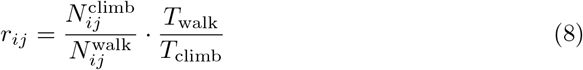

where *T*_walk_ and *T*_climb_ are the total times spent walking and climbing for all trials, respectively. The grand mean ratio 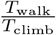was determined to be 0.563. If *r*_*ij*_ *>* 1 or *r*_*ij*_ *<* 1, the MP IN exhibited a bias for spiking during climbing or walking, respectively. The greater the deviation from 1, the more pronounced the bias.

#### Climbing classifier neuron (CCN)

In analogy to a binary classifier, the CCN was tuned to spike during climbing behaviour only, where ‘climbing’ defined as a body pitch angle exceeding 10° relative to the horizontal. To achieve this, the simplified LIF model proved adequate, with neuron and synaptic dynamics governed by Eqs (1), (2), and (3). A bias for ‘climbing behaviour’ was defined via a critical value *R*_ccn_ for Eq (8), whereas a bias for ‘walking behaviour’ was defined via its reciprocal value. Accordingly, primitive neurons with a significant climbing bias (*r*_*ij*_ *> R*_ccn_ *>* 1) were connected to the CCN through excitatory synapses with weight *ω*_ccn_, whereas those biased toward walking 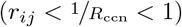 were connected through inhibitory synapses with the negative weight − *ω*_ccn_. Primitive neurons with a weak bias 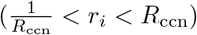 were not connected, i.e. *ω*_ccn_ = 0. The parameters *R*_ccn_ and *ω*_ccn_ were varied using a grid search to determine their optimal values.

To test the accuracy of the CCN, the experimentally measured body pitch time courses for each trial were segmented into 50 equally sized bins. For each bin, it was determined whether the CCN spiked (predicted condition) and whether the body pitch exceeded 10° (actual condition). Subsequently, a confusion matrix was constructed, with climbing designated as the positive condition and walking as the negative condition. The model’s accuracy was assessed using the MCC metric, determined through Eq (5), while varying *ω*_ccn_. The *r*_*ij*_ values were extracted from 20 trials separate from the trial used for optimization and estimation. This approach ensured that the model was ‘trained’ on climbing trials distinct from the ‘testing’ trial. The testing and training trials were rotated such that each trial served as the testing trial once, employing a method similar to leave-one-out cross-validation (LOOCV). The final MCC was calculated as the mean of all 15 LOOCV iterations.

#### Pitch estimator neurons (PENs)

As a complement for a binary classifier of climbing versus walking, the objective of the PENs was to encode body pitch through its spike rate over time. The concept for connectivity and tuning of both PENs was similar to that of the CCN, except that it included differential activity in two neurons. Climbing-biased MP (*r*_*ij*_ *> R*_pen_ *>* 1) were connected to *PEN*_+_ via excitatory synapses of strength *ω*_pen_, while walking-biased primitive neurons 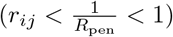 were connected to *PEN*_−_ through excitatory synapses (strength *ω*_pen_). The spike rate time course representing body pitch 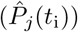 was obtained by subtracting the spike response of *PEN*_−_ from that of *PEN*_+_ with subsequent z-normalization. To assess the performance of the PENs, the experimental body pitch time course was interpolated to match the time step of the model (*dt* = 5 ms → 0.25 ms), and z-normalized (*P*_*j*_(*t*_i_)).

Again, the parameters *R*_pen_ and *ω*_pen_ were varied using a grid search to determine their optimal values. For optimisation, the squared difference between the z-normalized time courses was averaged for each discrete time step *i*, yielding the MSE_*j*_ for trial *j*:

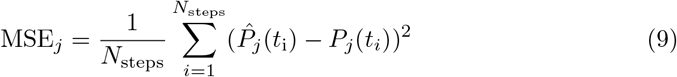

Similar to the procedure applied to the CCN, this process was repeated so that each trial served as the testing set exactly once, without being part of the training set. Finally, the LOOCV mean squared error (MSE) was computed using:

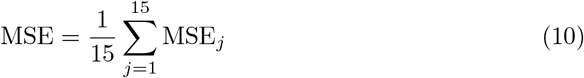

## Results

### Layer three: Movement primitive layer

With reference to behavioural and modelling results on insect locomotion, swing and stance phases of the step cycle are thought to involve distinct modes of leg movement control [34]. Whereas current models of adaptive locomotion use ground contact or load to switch between the mutually exclusive leg states swing and stance [35, 36], here we test whether an internal representation of these states may be based on kinematic parameters alone. To do so, we propose a layer of second-order INs that draws information from proprioceptive encoding of joint angles and velocities, as proposed in the companion paper (van der Veen et al., companion paper [26]). This movement primitive (MP) layer contained 672 MP INs that acted as coincidence detectors. We hypothesized that some of these second-order INs would be capable of encoding step cycle phases but also transitions between them. For each leg, 112 unique MP INs fired whenever two or three inputs overlapped with coinciding spikes. Each input was a spike train of a velocity or position IN from a different joint (*α, β, γ*) of a single leg. This section discusses the MP INs optimization and performance, analyses the encoding properties of the total population of second-order neurons, and examines the spike characteristics of the best-performing MP INs.

### Optimization and performance

In the LIF model (Eqs (1) to (3)), the parameters *τ* and *ω* were the only parameters subject to optimization. The time constant *τ* determined the time frame during which incoming spikes were considered as being temporally proximal. A relatively high *τ* value may have caused temporally distant spikes to trigger a postsynaptic response, potentially compromising accuracy. Conversely, a low *τ* value may have resulted in fewer overlapping presynaptic spikes within the designated time frame, leading to missing postsynaptic spikes. Therefore, optimizing the time constant *τ* was crucial to achieve optimal accuracy.

The synaptic weight *ω* determined the strength of a synapse connection. If the synaptic weight was too low, no postsynaptic spikes would occur in response to multiple presynaptic spikes. Conversely, if the weight was too high, subsequent spikes from a single input might trigger a response. Ideally, a spike from one input would set the membrane voltage close to the threshold, allowing a subsequent, temporally proximal spike to push the membrane voltage beyond threshold. Since ‘temporal proximity’ depended on *τ*, both parameters needed to be optimized together.

An additional consideration concerned the disparity in spike dynamics between the position and velocity INs. Position INs typically had a higher and more constant spike rate, while velocity INs exhibited a lower spike rate and activated and deactivated more rapidly. To take this into account, separate weights *ω*_pos_ and *ω*_vel_ were optimized for position and velocity IN spike train inputs, respectively. Moreover, some input combinations required distinct settings to perform optimally, especially ones with different numbers of inputs. When the weights that had been optimized for two inputs were applied to an IN with three inputs, the primitive neuron could spike with only two inputs overlapping, rather than all three. Consequently, the optimal weights for MP INs with three inputs were typically lower than those for two inputs. Therefore, *ω*_pos_ and *ω*_vel_ were optimized separately for each subset *E*_1_ through *E*_7_.

The synaptic weights of position INs (*ω*_pos_) and velocity INs (*ω*_vel_) were adjusted individually in a grid search. The weights ranged from 2 mV to 18 mV, with increments of 2 mV. Fig 2A shows how the weights for each subset depended on *τ*, where all weights correspond to the optimal MCC (Eq (5)). In general, the optimal weights decreased with increasing *τ*. As *τ* grew larger, the membrane voltage decayed more slowly and less synaptic strength was needed to reach the threshold potential. The p-v and p-p-v subsets were exceptions to this rule, with less consistent changes than in other subsets.

**Fig 2.**
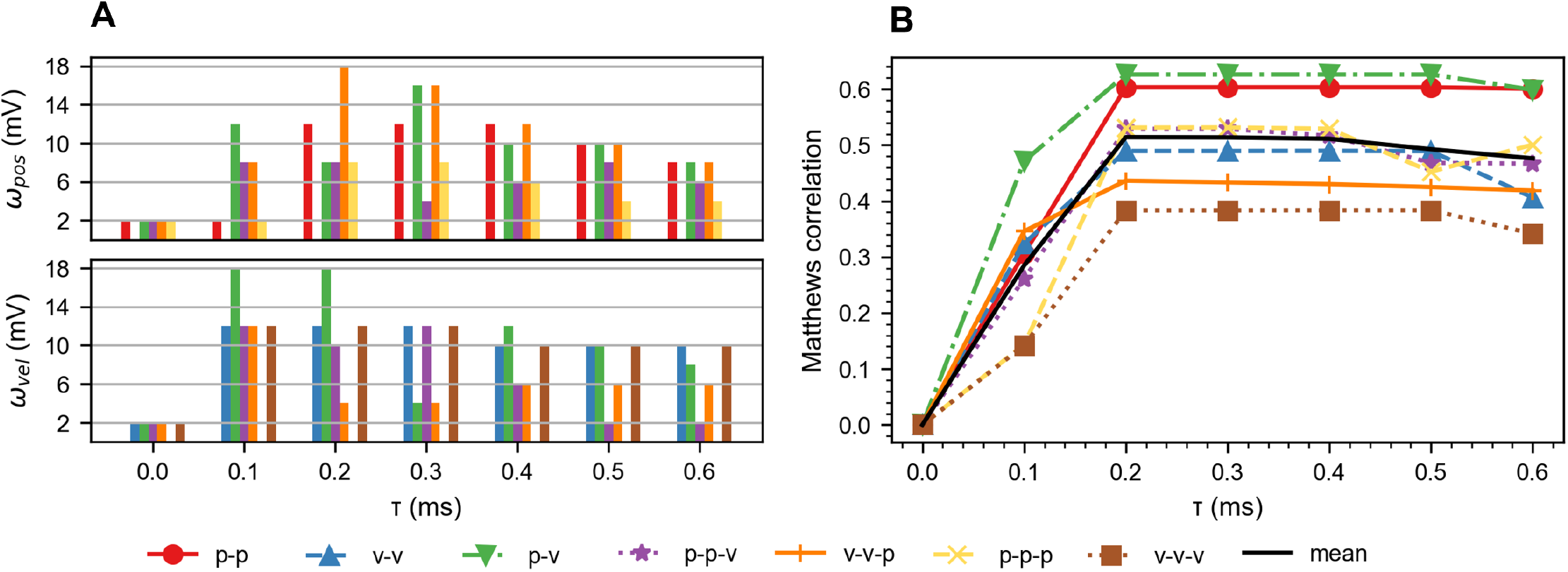
Optimization of synaptic weights and the time constant in the MP layer. **A**. Optimal synaptic weights for position inputs (*ω*_pos_, top) and velocity inputs (*ω*_vel_, bottom) optimized for seven subsets (*E*_1_ through *E*_7_) of MP IN for different time constants (*τ*). The synaptic weights ranged from 2 mV to 18 mV. They were adjusted to maximise MCC values, separately for each subset. **B**. Optimization criterion MCC as a function of *τ* for the seven subsets of MP IN, using the optimal weights from panel A. The black line shows the mean MCC, highlighting the overall performance of layer 3.

Fig 2B illustrates the MCC as a function of *τ* for all seven subsets of MP IN, using the optimal weights from Fig 2A. The performance peaked around *τ* = 0.2 ms and declined as the time constant increased further. There was a clear performance gap between subsets, with subsets comprising more position IN inputs (e.g. p-p) outperforming the subsets comprising more velocity IN inputs (e.g. v-v). Additionally, the subsets with two inputs (e.g. v-v) slightly outperformed those with three inputs (e.g. v-v-v).

#### Encoding properties

Next, we wanted to identify MP INs that encoded leg movement episodes that were characteristic for one of the alternating step cycle phases, i.e., stance or swing, or the transitions between them. Since the MP INs integrate information from all three joints of a leg, we hypothesized that some of these INs were phase-specific, firing either during swing or during stance, but not during the respective other phase. Similarly, we expected some MP INs to fire only during one of the transitions between these phases.

As the experimental dataset comprised annotations of all swing and stance phases of each leg, encoding of step cycle phases could be tested with a score that assessed the congruence of a spike train with the empirical step cycle time course. The match with corresponding templates of Eq (6) were scored with Eq (7). Since the swing and stance phases were mutually exclusive, a high score for the swing target vector indicated weak stance encoding. More specifically, a score of 1 indicated perfect swing encoding, while a score of 0 indicated perfect stance encoding. The same analogy applied to the transition scores: a score of 1 indicated perfect swing-to-stance encoding, while a score of 0 indicated perfect stance-to-swing encoding.

For illustration of this scoring procedure, Fig 3 shows the spike characteristics of a MP IN of the left hind leg (L3) with the input combination (*vel*_+_, *vel*_+_, *vel*_−_). This combination corresponded to simultaneous protraction at the *α* joint (top panel), levation at the *β* joint (2nd panel), and flexion at the *γ* joint (3rd panel). The activity of this particular MP IN achieved the highest swing score among all 672 third-order INs (4th panel in Fig 3). Note that this MP IN fired only if spikes from all three inputs coincided, yielding a nearly perfect overlap with the annotated swing state (value zero) of the experimental data (bottom panel). In fact, the temporal overlap of value zero in panel 5 (‘swing’) and spike episodes in panel 4 is considerably better than with spike episodes of any of the three single inputs (panels 1 to 3). To illustrate the consistency of this neuron’s response, Fig 4A shows its likelihood to spike in any one of the 16 bins of the standard step cycle. As for the single trial example of Fig 3, the neuron shows consistent swing encoding, with a near 100 % likelihood to spike during swing phase and 0 % during stance, with smooth transitions occurring at the onset and offset of swing.

**Fig 3.**
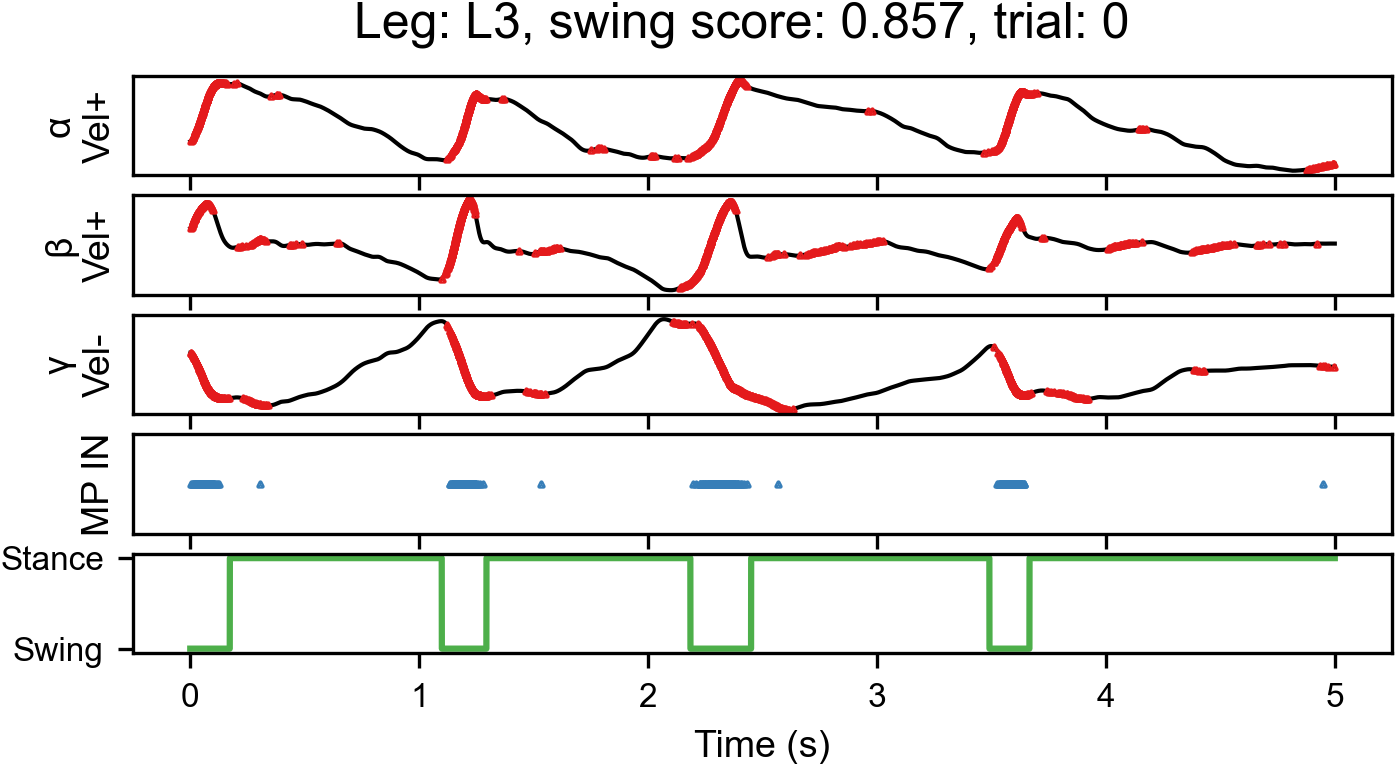
Presynaptic and postsynaptic spike trains of the best-performing swing encoder. The top three panels show the joint angle traces of the *α, β*, and *γ* joints of the left hind leg (L3). Red dots superimposed on the joint angle time courses mark single spike events of the presynaptic input IN in layer 2 for each joint. For instance, the *vel*_+_ neuron of the *α* joint spikes during protraction at the thorax-coxa joint. The spike trains shown in the top three panels served as inputs to the best-performing swing encoder of the MP layer (panel 4). This IN spiked whenever all three inputs spiked coincidentally. The bottom panel illustrates the annotated stance (high) and swing (low) phases of the leg movement of this experimental trial. Note that the timing of neural activity in this v-v-v IN of layer 3 matched the leg’s swing phase much better than any of its inputs from velocity IN layer 2.

**Fig 4.**
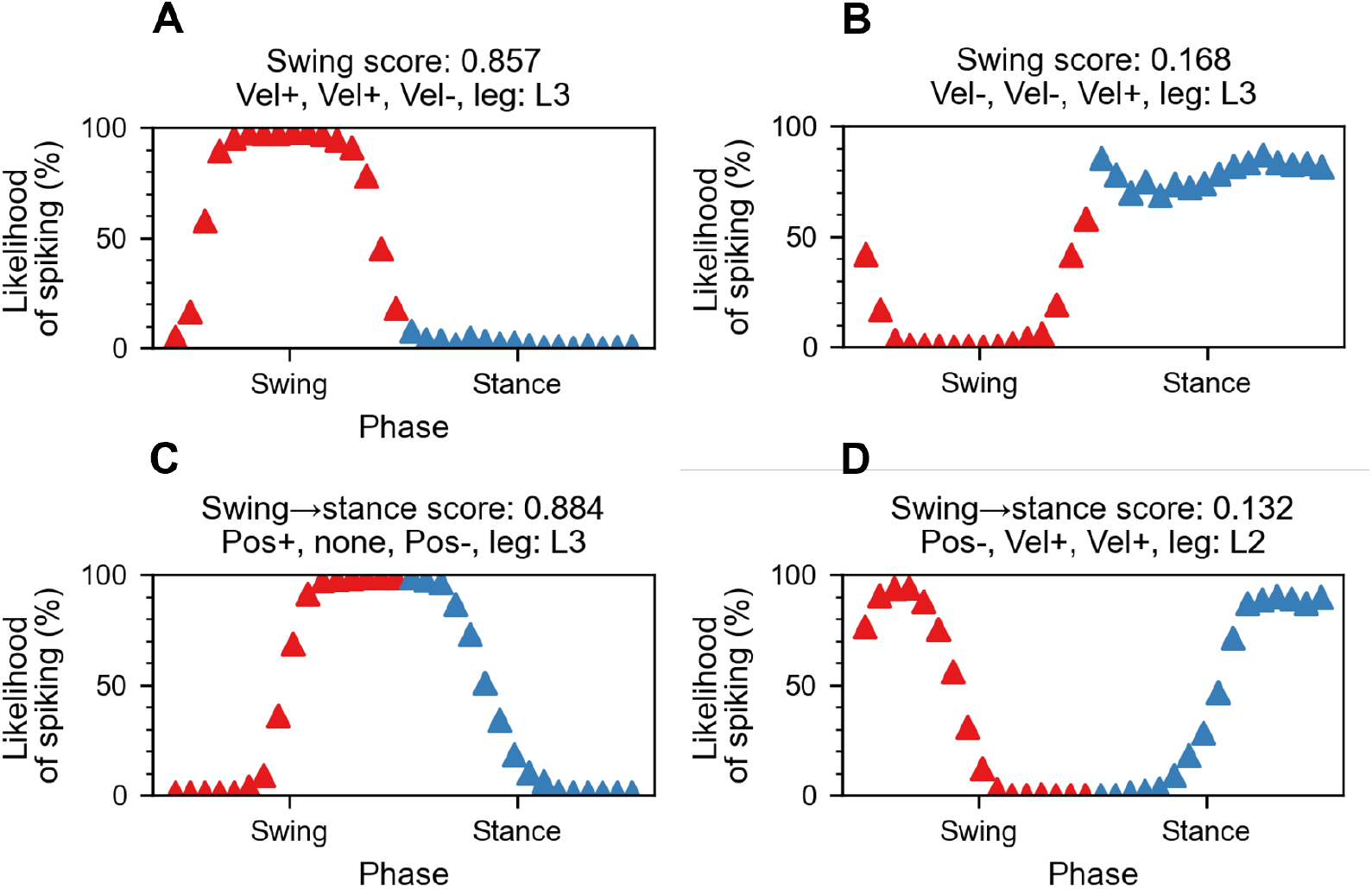
Likelihood of neuronal spiking during the swing and stance phases of locomotion. Each panel displays spike likelihood of the highest-performing INs among all 672 MP across a standard step cycle. Each panel corresponds to a specific step cycle phase or phase transition: **A**. swing, **B**. stance, **C**. swing-to-stance, and **D**. stance-to-swing (for templates see Eq (6)). The x-axis represents the phase of the step cycle, divided into swing and stance periods and further separated into 8 equally sized bins. The y-axis shows the likelihood of that neuron to spike in a particular time bin, expressed as a percentage. Red and blue triangles label swing and stance phase, respectively. The corresponding scores and input combinations are given in the panel headlines. Note that the best-performing INs of the MP layer belonged to the v-v-v subset in case of swing/stance encoding, to the p-p subset for swing-to-stance transitions, and to the p-v-v subset for stance-to-swing transitions.

In analogy, the remaining panels of Fig 4 show the performances of the best-encoding MP INs for stance (Fig 4B), the swing-to-stance transition (touch-down: Fig 4C), and the stance-to-swing transition (lift-off: Fig 4D).

Fig 5 was constructed to determine which first-order INs contributed most to swing and stance encoding. To achieve this, the swing scores for all MP IN were grouped according to velocity and position IN inputs for the three joints and six legs. The statistical summary of Fig 2 is provided in Table 2, which tabulates the scores of paired t-tests for all *vel*_−_-*vel*_+_ and *pos*_−_-*pos*_+_ couples per joint and leg. Significance levels are colour-coded in red (p*<*0.1 %), blue (p*<*1 %), and green (p*<*5 %). High t scores were expected for second layer INs that fired more often during a particular phase relative to their negative counterpart, e.g. *vel*_+_ and *vel*_−_ in the *α* joints.

**Table 1.** Model type and parameter values for the second- and third-order INs after optimization.

| IN | Type | $E_L$ | $V_T$ | $\tau$ | $\omega$ | R |
| --- | --- | --- | --- | --- | --- | --- |
| MP | LIF | -70 mV | -50 mV | 0.2 mV | see Fig 2A | - |
| CCN | LIF | -70 mV | -50 mV | - | 0.15 mV | 1.8 |
| PEN | LIF | -70 mV | -50 mV | - | 10 mV | 1.62 |

**Table 2.** Comparison of swing scores between + and - INs. Summary of two-sided, paired t-tests comparing swing scores of *vel*_−_ vs. *vel*_+_ and *pos*_−_ vs. *pos*_+_ in Fig 5 across six legs and three joints. For sample sizes of *n* = 24 pairs per joint, the critical values were 3.792 for *p <* 0.001 (red), 2.819 for *p <* 0.01 (blue), and 2.074 for *p <* 0.05 (green). Black numbers indicate non-significant results. Positive t-scores indicate stronger swing encoding associated with *vel*_+_ or *pos*_+_, while negative t-scores indicate stronger swing encoding associated with *vel*_−_ or *pos*_−_.

| INs | joint | R1 | R2 | R3 | L1 | L2 | L3 |
| --- | --- | --- | --- | --- | --- | --- | --- |
| $vel_+$<br>&<br>$vel_-$ | $\alpha$ | 11.75 | 14.47 | 10.27 | 11.51 | 14.57 | 10.7 |
| | $\beta$ | -2.99 | 2.03 | 6.78 | -4.14 | 2.37 | 7.34 |
| | $\gamma$ | 10.13 | -4.86 | -10.0 | 11.12 | 0.79 | -9.44 |
| $pos_+$<br>&<br>$pos_-$ | $\alpha$ | -0.95 | -4.1 | -5.23 | -0.19 | -2.51 | -4.63 |
| | $\beta$ | 3.36 | -7.07 | -4.47 | -0.29 | -6.86 | -7.0 |
| | $\gamma$ | -9.01 | -10.0 | -2.92 | -8.44 | -9.58 | -3.33 |

**Fig 5.**
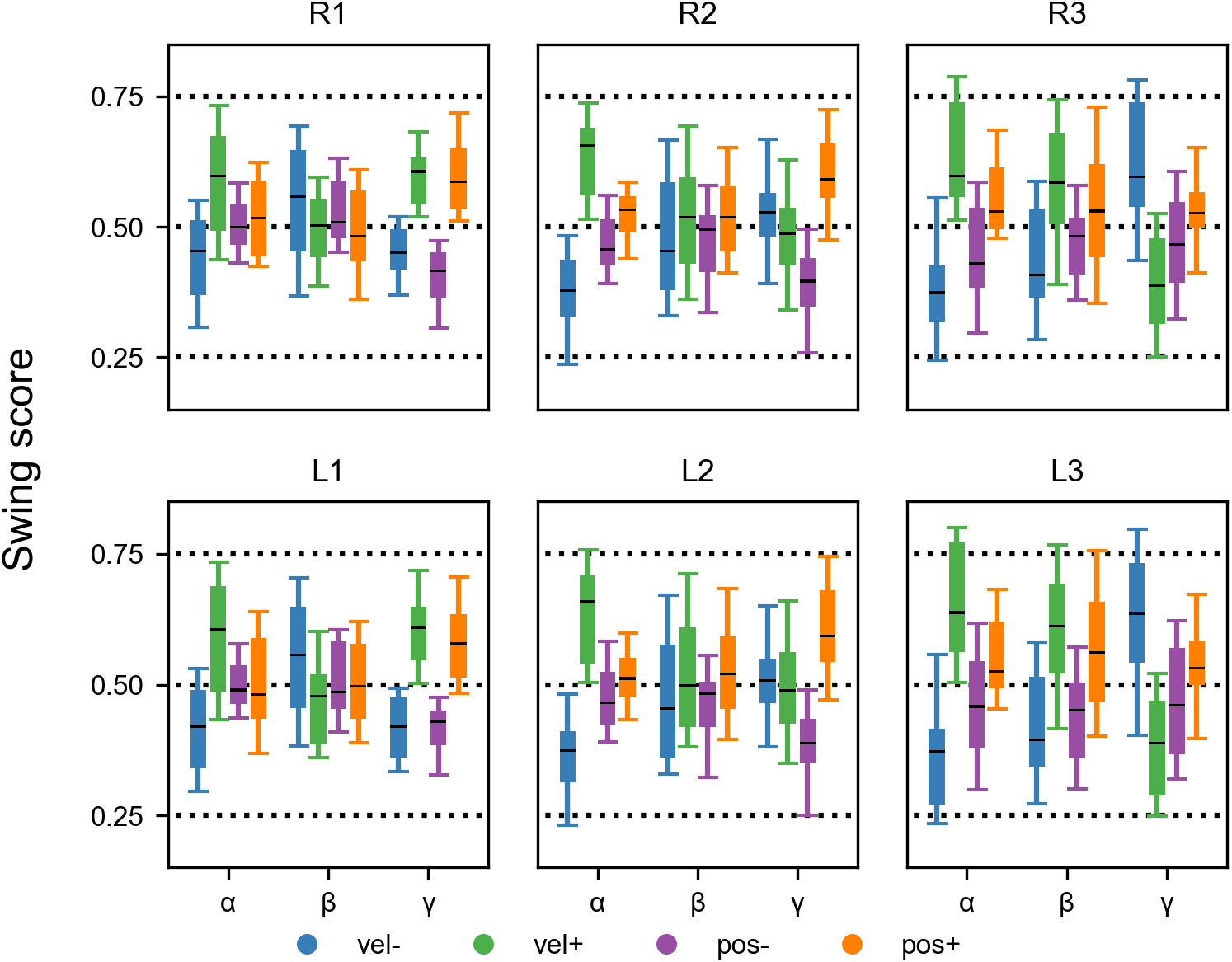
Joint- and leg-specific contributions to the encoding of swing and stance. Box plots group swing scores of all MP IN according to their to velocity (blue and green) or position (purple and orange) inputs. The six panels group information from one leg, with 3 *×* 4 box plots grouping the information from individual joints. In each box plot the black line represents the median value, the box encompasses the inter-quartile range, and the whiskers comprise the central 90 % of the data. For statistics associated with this figure, see Table 2.

Overall, the result was highly symmetrical for left and right legs (compare top and bottom panel rows in Fig 5) with a slight asymmetry occurring in velocity inputs of the *β* and *γ* joints of the middle legs (compare columns R2 and L2 in Table 2). For several joints, Fig 5 reveals a functional reversal between front and hind legs. As an example, the *vel*_+_ inputs of the *γ* joint contributed to swing in the front legs but to stance in the hind legs, with the opposite being true for *vel*_−_ inputs of the *γ* joint (compare right-most green and blue box plots in R1/L1 panels versus in R3/L3 panels). Similarly, though with opposite tuning, *vel*_+_ and *vel*_−_ inputs of the *β* joint swapped from encoding of swing and stance between front and hind legs. These front-to-rear changes are reflected by sign reversals of the t-values in Table 2.

A notable exception to this kind of front-to-rear change concerns the *vel*_+_ and *vel*_−_ INs of the *α* joint that showed consistent biases towards swing or stance encoding, respectively, in all legs (see left-most green and blue box plots per panel). This result reflects the fact that in forward walking animals, the *α* joints tend to be protracted during swing and retracted during stance. Front legs differed from middle and hind legs in that there was little or no difference between the swing scores of *pos*_+_ and *pos*_−_ inputs for both the *α* and the *β* joint (see low t-scores in Table 2).

To summarize the overall tuning of MP INs, the histograms in Fig 6 depict the distribution of scores for two target vectors: the swing phase and the swing-to-stance transition. The histograms are grouped by front, middle, and hind legs, with each group comprised 224 MP INs. Neurons with scores above 0.75 and below 0.25 are highlighted as ‘strongly encoding INs’. Additionally, data-driven thresholds were defined as below the 5th percentile and above the 95th percentile. The numbers of strongly encoding INs show that MP INs of the middle and hind legs were more effective as phase encoders compared to those of the front legs. This is also reflected in the overall width of the histograms, i.e. the largest 90% percentile range. For phase transitions, more MP INs exceeded the threshold than for swing/stance encoding. Phase transition encoding was strongest for the hind legs, moderate for the middle legs, and extremely weak for the front legs. For detailed information about the most efficient input combinations, Supplementary Tables S2 Table and S3 Table list the input combinations of the six highest-scoring MP INs for the middle and hind legs, respectively. These tables underscore the relevance of *vel*_+_ and *vel*_−_ INs of the *α* joint for reliable swing/stance encoding. In contrast, *pos*_+_ and *pos*_−_ INs of the *α* joint proved to be relevant for encoding of phase transitions.

**Fig 6.**
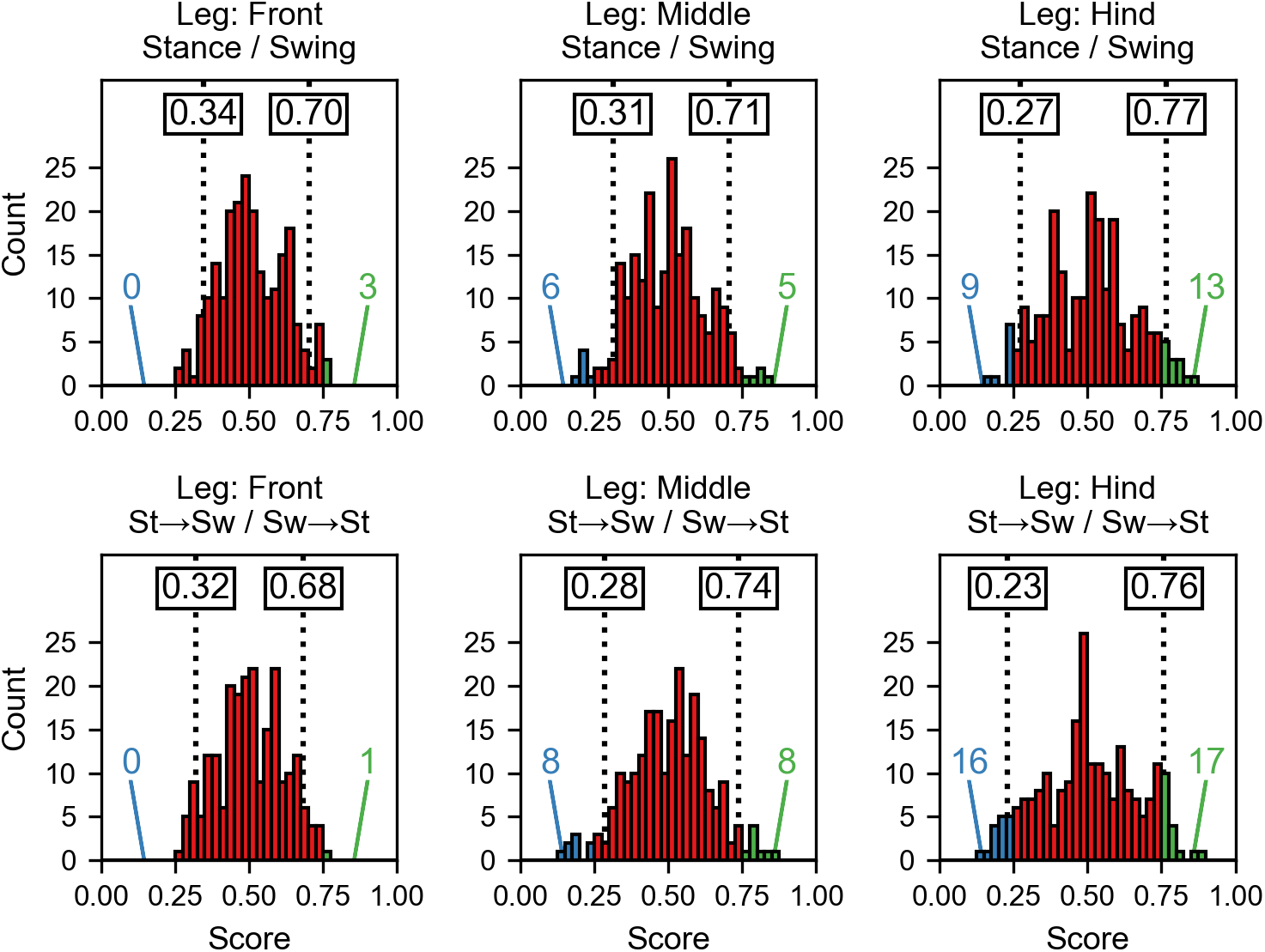
Histograms of swing/stance and transition scores for different leg types. Each histogram displays the distribution of encoding scores of 224 INs. Pooled for left and right legs, the top row shows swing/stance encoding with separate panels per leg. A score of one indicates a perfect swing encoder, while a score of zero indicates a perfect stance encoder. The second row presents the scores for the encoding of phase transitions, where a score of one indicates a perfect swing-to-stance encoder and a score of zero indicates a perfect stance-to-swing encoder. Blue and green numbers at the bottom of each histogram report the counts of strongly encoding INs, with scores below 0.25 or above 0.75, respectively. The numbers inside the boxes on the dotted lines label the lowest 5 % and highest 5 % of the distribution, respectively.

### Layer four: Posture layer

The fourth and final layer of our model was developed to test how well distributed proprioceptive information about whole-body movement may be read out to encode high-level parameters of locomotion. Given that our experimental data comprised repetitive climbing intervals, we decided to focus on the pitch angle of the body axis. Body pitch is particularly suitable to test proprioceptive encoding of high-level parameters in general, because (i.) there is no sensory organ dedicated to measuring body pitch, (ii.) it is highly dependent on leg postures, though (iii.) the relative contributions of the six legs change all the time because of alternating ground contact. Therefore the main objective for fourth layer was to devise a ‘posture layer’ suitable to estimate the pitch angle of the metathorax from kinematic information from 18 leg joints. To this end, the posture layer comprised a total of three third-order INs: one being a binary classifier tasked with the classification of climbing versus walking (CCN), and the other two being responsible for providing a continuous body pitch time course (PENs).

In the experimental trials, the setup required transient, large-amplitude adjustments of the stick insects’ body pitch due to high physical stairs (see Fig 1b). The height of these stairs was about three times the body clearance and exceeded leg length. As a consequence the stick insect alternated between regular walking and climbing (see Fig 1ai). During the walking phase, the insect’s body pitch remained close to zero with minor fluctuation, indicating that the thorax was nearly parallel to the substrate. In the climbing phase, the body pitch exceeded 10°, reaching up to approximately 60°, as illustrated in Fig 1b.

#### Climbing classifier neuron (CCN)

Given that level walking and climbing were easily distinguishable by means of a simple threshold, we used an arbitrary body pitch angle of 10° to tell climbing from walking. Accordingly, the CCN was devised as a binary classifier neuron that was to spike whenever the body pitch angle exceeded 10°. Classification was optimized by maximizing the MCC (Eq (8)) for binarized experimental body pitch time courses with a threshold of 10°. The parameters to be tuned were the critical value *R*_ccn_ that selected climbing-sensitive MP INs (Eq (5)), and their synaptic weight *ω*_ccn_.

Fig 7A plots the MCC while varying *ω*_ccn_ and *R*_ccn_ in a grid search. The optimal values were *ω*_ccn_ = 0.15 mV and *R*_ccn_ = 1.8, reaching an average MCC of 0.597. At this ratio, the middle legs contributed the most information to posture estimation, with 46 connected neurons. The front legs contribute the second most, with 35 connected neurons, while the hind legs contributed the least, with only 28 connected neurons. Fig 7B illustrates a single LOOCV iteration that scored closest to the mean MCC value, representing the average performance. Spikes mainly occured when body pitch exceeded 10°, while some false positives occurred when the body pitch was below 10°.

**Fig 7.**
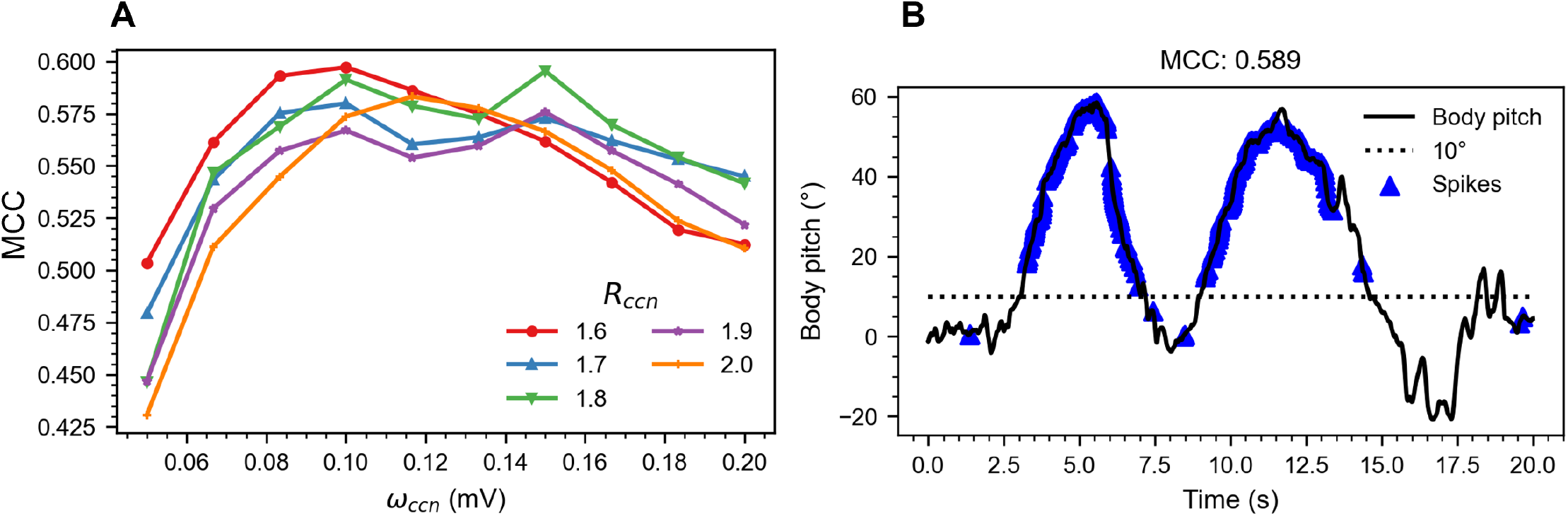
Optimization and performance of the binary pitch classifier neuron CCN. **A**. Mean MCC of 15 LOOCV iterations for the CCN while varying *ω*_ccn_ and *R*_ccn_. **B**. A single LOOCV iteration that scores closest to the mean LOOCV values plotted in panel A, i.e., for optimal values *ω*_ccn_ = 0.15 mV and *R*_ccn_ = 1.8. The CCN mainly spiked when body pitch exceeded 10°.

#### Pitch estimator neurons (PENs)

As a complement to the CCN, two pitch estimator neurons (PENs) were tuned to encode the body pitch angle by continuous variation of their spike rate. As for the CCN, this required optimization of a walk/climb ratio (*R*_pen_) for presynaptic MP INs, and a corresponding synaptic weight *ω*_pen_. Fig 8A shows the MSE, calculated using Eq (9), as a function of *ω*_pen_ and *R*_pen_. The optimal parameters were found to be *ω*_pen_ = 10 mV and *R*_pen_ = 1.620, resulting in an MSE of 0.664. Performance improved (lower MSE) with increasing synaptic weight until a plateau was reached, suggesting that the model was optimal when all presynaptic spikes from the selected MPs INs triggered a postsynaptic spike. At these optimal synaptic weights, the PENs functioned as perfect spike integrators.

**Fig 8.**
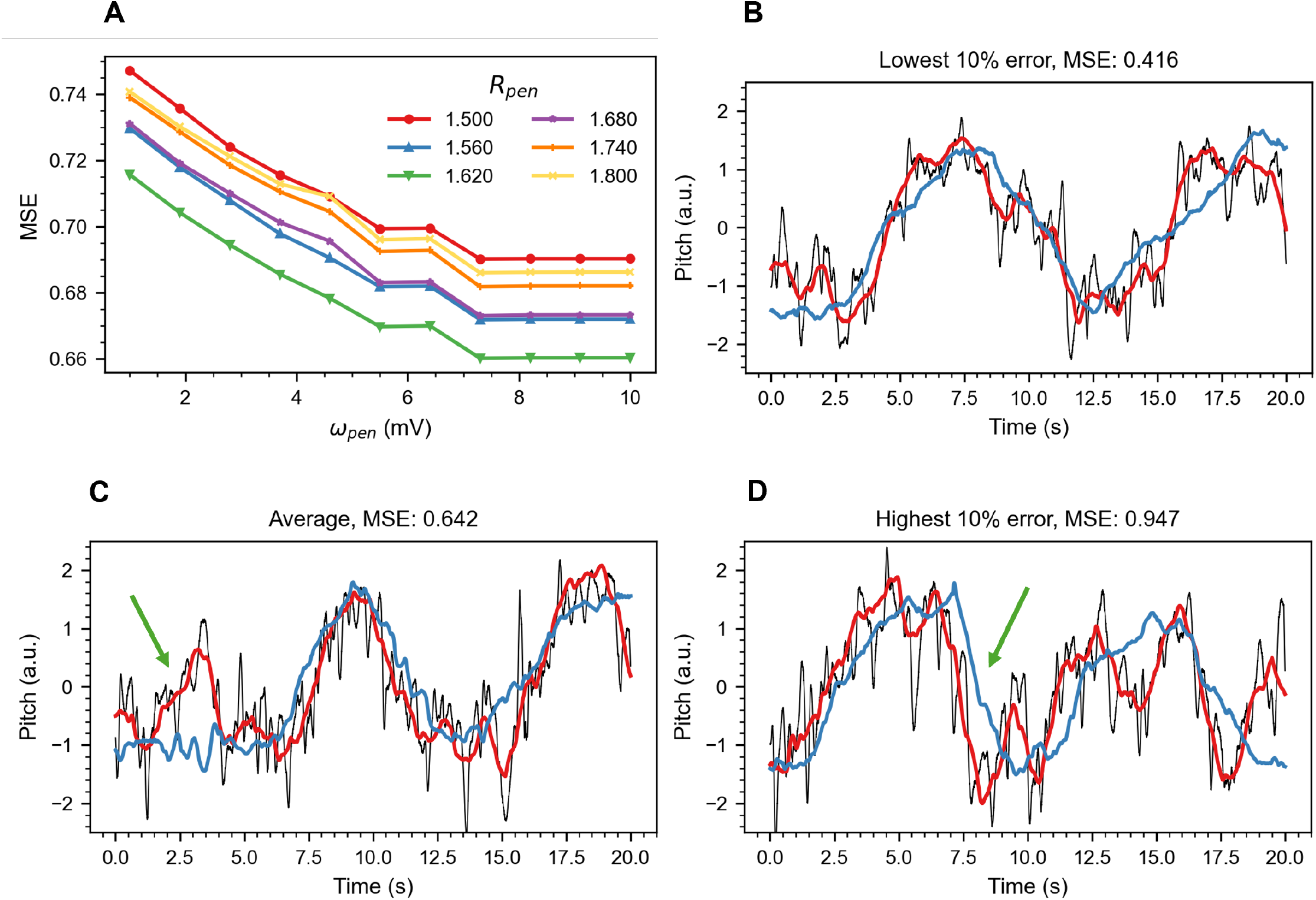
Optimization and performance of posture-estimating neurons (PENs) **A**. Mean MCC of 15 LOOCV iterations for the PENs while varying *ω*_pen_ and *R*_pen_. **B** to **D** show single iterations for optimal values *ω*_pen_ = 7.0 mV and *R*_pen_ = 1.62, selected according to their performance. The blue line represents the ground truth, the black line shows the model prediction, and the red line indicates a moving average of the model prediction. **B**. Top 10 % performing LOOCV iteration. **C**. Iteration closest to the mean LOOCV value. Notice the incorrect prediction of an entire peak, indicated by the green arrow. **D**. Bottom 10 % performing iteration. Notice that the estimate leads the ground truth (green arrow).

Fig 8C illustrates tuned PEN performance, using the single iteration with its MCC closest to the mean, representing the average performance. The response of the PEN aligned crudely with experimental data, showing significant noise. For better comparison with the ground truth, the red line shows the running average of the PENs spike rate. The alignment with the ground truth was good for most of the trial, though a brief episode clearly deviated with a conspicuous peak (see green arrow in Fig 8C). For comparison, the range between the 10 % best- and least-performing scenarios are depicted in Fig 8B and Fig 8D, respectively, with 80 % of all iterations falling within these two performances. Whereas deviation from the ground truth was small in Fig 8B, there was a conspicuous lead in some episodes of 8D (see green arrow).

## Discussion

We conclude that distributed proprioception of joint angle and joint angle velocity at three joints per leg is sufficient for reliable encoding of the step cycle phases swing and stance, as well as transitions between these phases (Figs 3, 4 and 6). We attribute this to pronounced swing- or stance-specific leg postures and movements at several leg joints (Fig 5). The pool of single-leg movement primitive INs of layer three was sufficient to supply proprioceptive information about whole-body kinematics. This was demonstrated by binary classification of climbing versus walking (Fig 7), as well as by spike-rate encoding of the high-level movement parameter body pitch (Fig 8).

### Layer three: Movement primitive layer

The third layer was termed the ‘movement primitive layer’, inspired by the hypothesis that complex movements are composed of elementary motor synergies [31, 54] that arise from concurrent activity of multiple muscles. Motor synergies are thought to underlie elementary patterns of coordinated movement of multiple joints, called movement primitives [32]. This concept has been studied for motor behaviours of both invertebrates and vertebrates, with evidence at behavioral, muscular, and neural levels. For example, human reaching movements can be described by small sets of coordinated movement patterns associated with couplings of multiple degrees of freedom [55]. Movement primitives are also applied in the development of humanoid robots [56].

Following the movement primitive concept, the first objective of this paper was to construct a layer of second-order INs that coupled position or motion information across two or three joints. Since the leg motorneurons of insects are organised in 3 *×* 2 thoracic hemiganglia [57], i.e. one per leg, our movement primitive INs received input from two or three joints per leg. Accordingly, their activity signalled a particular pattern of intra-leg coordination, independent of the state of their neighbouring legs. Owing to the distinct spike activity of their presynaptic neurons, MP subsets with input from position INs achieved higher accuracy compared to those receiving input from velocity INs (see Fig 2B). This discrepancy could be attributed to three factors: First, the spike rate response of velocity INs was delayed by approximately 25 ms relative to the ground truth, resulting in errors in the MP layer. Second, the mean spike rate of position IN was higher and sustained for extended periods when joint angles were high. In contrast, velocity INs exhibited shorter spike peaks, leading to a lower total spike count than in position INs. Lastly, position INs maintained sensitivity across the entire receptive field of a tactile hair, unlike velocity INs, whose sensitivity was closely linked to the number of hairs in a hair plate and only fires during the onset of a complete hair deflection (van der Veen et al., companion paper [26]). Additionally, a performance gap was observed between subsets with two inputs as opposed to three inputs. Generally, subsets with two inputs outperformed those with three inputs. This discrepancy arose from the fact that three inputs were less likely to spike coincidentally than two inputs, consequently reducing the MCC.

Since our MP INs layer comprised all 112 possible combinations of position- and motion inputs from one leg, we wanted to evaluate the behavioural relevance of their tuning properties on the basis of experimental data. We did so in two steps, using annotated data from unrestrained walking stick insects [47]. In a first step, we compared the relative differences of the three leg pairs, assuming bilateral symmetry of encoding properties for left and right legs. Indeed, the top and bottom panels of Fig 5 show nearly the same results for right and left legs, respectively (with largest differences for *γ* velocity encoding in middle legs; Table 2). In contrast, Fig 5 reveals distinct front-to-rear differences in the swing/stance specificity of particular leg joints, both for position- and velocity-encoding INs. This reflects different functions of the three leg types [58], including their inherently different walking directions [59]. This is in line with our finding in Fig 6 that MP IN of the front leg performed worse as swing, stance, or transition encoders, which can be attributed to the front legs’ more irregular stepping pattern [60, 61], engagement in searching behavior [62], and more short steps [47]. Increased irregularity of front leg stepping would have caused MP INs to activate less consistently throughout step cycle, potentially reflecting mixtures of swing and stance phases, strongly reducing the encoding performance for particular step cycle phases. In contrast, the locomotion cycle of the middle and hind legs is more regular, resulting in more movement primitives being associated with a specific phase. Consequently, a greater number of MP IN encode for swing, stance, or transitions, as illustrated by the broader distribution in Fig 6.

In a second step, we looked at specific encoding properties of particular MP INs. Similar to the examples shown in (Fig 4A and B), INs observed in locusts or cats [37, 38] have been shown to be active predominantly during either the swing or stance phase of the step cycle. These INs may be considered as internal representations of swing and stance phases for each leg, which have been proposed to reflect distinct control modes of the step cycle [34], owing to the distinct boundary conditions due to mechanical coupling through the substrate. Fig 4C and D depict MP INs whose spike activity signal transitions between step cycle phases. Experimental results show that these transitions are important events in the control of the step cycle [63], but also for inter-leg coordination. For example, the swing-to-stance transition, i.e., touch-down of a leg is an important event in the timing of the lift-off in the next anterior leg coordination [40, 41]. Similarly, the end of the stance phase and the transition to swing phase affects timing events in neighbouring legs [64, 65]. Indeed, this transition point information may be signalled by the coxal hair plates [66]. Several biomimetic robots and locomotion models explicitly include distinct swing and stance modules [35, 36, 67, 68]. Our results suggest not only a proprioception-based internal representation of the leg’s state (swing/stance) but also a more detailed encoding of different events throughout the step cycle. Furthermore, the results demonstrate that this representation can be derived from limb kinematics alone, that is, without information about load or body-substrate interaction. Although this is consistent with the finding that limb kinematics changes only little when walking on slopes, despite drastic changes in load [69], future studies will need to test whether step-cycle specific neural activity may be maintained with kinematic proprioceptive information, only.

### Layer four: Posture layer

The second objective of this paper was to test to what extent high-level information about body posture could be derived from distributed, low-level proprioceptive information. Visual neuroscience has established a number of examples, where high-level information about general locomotion parameters such as flight velocity [70] or distance travelled [71] are derived from large field optic flow sensed by a grid of local motion detectors. The mammalian visual system has become a textbook example for the representation of increasingly complex visual features [72]. More generally, cell assemblies of the mammalian cortex are commonly viewed to build a distributed neural representation of behaviorally relevant entities [73]. Assuming that the properties of the MP layer described above provided a distributed representation of the current locomotor state of the animal, we hypothesised that that the activity pattern in this layer should allow us to read out high-level information about the body posture and movement.

Indeed, an earlier study showed that an Artificial Neural Network could be trained to extract the body pith angle from distributed spike activity patterns of proprioeptive hair field afferents [45]. In taking motivation from this result, the present paper aimed to extract the same high-level parameter with a hierarchical spiking neural network model. Like the earlier study, we used the inclination of the metathorax relative to the horizontal as a measure of body pitch (Fig 1aii). While this angle is largely determined by the posture of the middle and hind legs with ground contact, the subset of these four legs having ground contact changes all the time. Moreover, the neurons of the first three layers of our network were optimized using experimental data without climbing episodes, i.e., from level walking trials only. In contrast, the third-order INs of the posture layer were evaluated with data from experimental trials with two climbing intervals, including proprioceptive information from the front legs, despite the fact that their contribution to metathorax inclination was much less direct than that from middle and hind legs.

The overall ability of the climbing classifier neuron to tell ‘climbing’ from ‘walking’ proved to be very good, though its performance measure was reduced significantly by occasional spikes that fired at the wrong time (Fig 6B). During periods of high body pitch, the number of spikes increased rapidly, rendering this classifier a reliable indicator for climbing in the stick insect. However, a similar result could have been obtained from attaching a high-pass filter to the body pitch estimator neurons.

The PENs estimated body pitch with an MSE of 0.660. The middle legs were found to be the most relevant for body pitch estimation, closely followed by the front legs, with the hind legs being the least relevant. This was somewhat different from the result of Gollin and Dürr [45], who also found the middle legs to be the most relevant, however followed by the hind legs and the front legs being least effective. We attribute the different result to the information processing of our first- and second-order interneurons, which was lacking in [45]. For example, suitable MP INs may have signalled a strongly raised front legs, which is a posture that could be indicative of spatial searching immediately prior to climbing. Considerations along these lines could also explain the apparent lead of some PEN solutions (e.g., Fig 8D), if body postural adjustments immediately prior to climbing may contribute to overestimation of the current body pitch angle. Overall, Fig 8 demonstrates that body pitch estimation is achievable with a hierarchical spiking neural network, reinforcing the idea that neural circuits in the stick insect can represent body pitch from distributed proprioceptive cues. The remarkable accuracy of this result is noteworthy, considering that the proprioceptive information passed through three spiking layers, compounding errors along the way. Future work may take into account that selecting presynaptic input elements based on their spike biases may not be particularly efficient. Instead, optimizing the synaptic weights through learning algorithms might improve the results even further.

### Future scope

We have identified four aspects of the proposed model, where future studies could further improve functionality and neurobiological plausibility: (i) the presence of noise, (ii) multimodal integration, (iii) encoding of further Degrees of Freedom, and (iv) spatial coordination. Concerning the first of these, a major computational difference between our SNN model and an animal’s CNS is that our model is deterministic, whereas real neural networks are not. Noise is an inherent aspect of all biological neuronal systems, present at every level of the CNS, from sensory to motor levels [74]. A future work may explore how noise originating from intrinsic stochastic properties of spike generation or synaptic transmission could enhance the biological accuracy of the computational model.

A second point concerns the fact that our model receives information from one type of proprioceptor only, and the corresponding proprioceptor model is tuned with a single set parameters. While the properties of peripheral proprioceptive encoding are discussed in more detail in the companion paper [26], the choice of proprioceptor type and its tuning properties must also have substantial effects on internal representations of movement primitives and whole-body posture. One reason is that the sensitivity of hair field afferents is limited, for example concerning the detection threshold for movement. As a consequence, including proprioceptors with higher movement sensitivity (e.g., chordotonal organs; [75]) could improve accuracy of velocity estimation. More generally, different insect proprioceptor types encode distinct physical magnitudes [24], allowing for multimodal encoding of kinematic variables such as position and motion along with dynamic variables such as stress or strain. In stick insects, afferent information from distinct proprioceptor types converges at various levels of proprioceptive encoding [76, 77] calling for integration of multimodal proprioceptive encoding in future computational models [78].

A third aspect concerns the encoding of additional degrees of freedom of motion, other than body pitch. Whereas, in principle, our model should be able to encode other high-level parameters such as speed or yaw rotation, future studies are needed to provide suitable experimental data to be used as ground truth. Potential expansions of our SNN may include additional third-order neurons to encode all three degrees of freedom of whole-body turning, i.e., yaw, pitch and roll, along with parameters associated with translational movement, such as forward speed or body clearance. Corresponding experimental data would have to link leg kinematics (or dynamics) to body axis position and orientation during natural turning, e.g., by linking 6 *×* 3 joint angle time courses to yaw-pitch-roll rotational velocities as the animal maneuvers around a pole.

Finally, an important aspect of motor flexibility in natural locomotion concerns spatial coordination among limbs [34], including spatial coordination of footfall patterns [10, 11] and targeted limb movements toward the body surface [7, 8, 16] or external objects [3–5]. Conceptually, this concerns the encoding of spatial coordinates within the immediate environment of the animal’s body [14] and sensorimotor transformations [30], including the coordinate transfer among limbs [13, 14]. To some extent, these concepts could be included into our SNN by expanding the movement primitive layer by neurons that signal coordinated action of neighbouring legs. Another possibility could be to use MP INs to indicate the optimal timing for a leg to transmit information to its rear neighbour, thus selecting the appropriate target position to be used for aimed foot contact. Such mechanisms of inter-leg coordination could improve the SNN’s potential to account for natural climbing behaviour beyond the encoding of body pitch.

## Supporting information

**S1 Fig LIF Dynamics**. The LIF model responds to a constant spike rate (333.3 Hz), governed by Eqs (1, 2, 3). A presynaptic spike, Pre, increases the membrane potential by 10 mV, and if no spikes are present, the membrane voltage, *V*, decays back to *E*_L_ = −70 mV. When *V > V*_T_ = −50 mV, a postsynaptic spike, Post, is recorded, and *V* resets to *E*_L_ = − 70 mV. The postsynaptic spike rate remains constant in response to a constant presynaptic spike rate, since the LIF model has no adaptation. At the shown input spike rate, every third presynaptic spike triggers a postsynaptic spike. The model parameters are given in (van der Veen et al., companion paper [26]), their Table 1 (velocity neuron).

**S2 Table Top 6 highest performing MP IN for the middle legs, with respect to phase and transition encoding**. The first term corresponds to the kinematic IN of the *α* joint, the second to the *β* joint, and the third to the *γ* joint. Some MP IN repeat since there is one for each side.

**S3 Table Top 6 highest performing MP IN for the hind legs, with respect to phase and transition encoding**. The first term corresponds to the kinematic IN of the *α* joint, the second to the *β* joint, and the third to the *γ* joint. Some MP IN repeat since there is one for each side.

## Acknowledgments

The authors would like to acknowledge the financial support of the CogniGron research center and the Ubbo Emmius Funds (Univ. of Groningen). And the authors would like to thank Arne Gollin for supporting the project by sharing a curated motion capture dataset and for providing us with Fig 1a,b,c.

